# An obligate symbiont of *Haematomyzus elephantis* with a strongly reduced genome resembles symbiotic bacteria in sucking lice

**DOI:** 10.1101/2024.11.14.623662

**Authors:** Jana Martin Říhová, Roman Vodička, Václav Hypša

## Abstract

The parvorder Rhynchophthirina with a single genus *Haematomyzus* is a small group of ectoparasites related to sucking and chewing lice. Previous screening based on the 16S rRNA gene indicated that *Haematomyzus* harbour a symbiotic bacterium whose DNA exhibits a strong shift in nucleotide composition typical of obligate mutualistic symbionts in insects. Within Phthiraptera, the most dramatically reduced genomes are found in the symbionts associated with sucking lice, living exclusively on mammal blood, compared to the less modified symbionts inhabiting the chewing lice, which feed on skin derivates. In this study, we investigate the genome characteristics of the symbiont associated with *Haematomyzus elephantis*. We sequenced and assembled the *Haematomyzus elephantis* metagenome, extracted a genome draft of its symbiotic bacterium, and show that the symbiont has a significantly reduced genome, which is with 0.39 Mbp the smallest genome among the symbionts known from Phthiraptera. Multigenic phylogenetic analysis places the symbiont into one of three clusters composed of long-branched symbionts from other insects. More specifically, it clusters together with symbionts from several other sucking lice, and also with *Wigglesworthia glossinidia*, an obligate symbiont of tsetse flies. Consistent with the dramatic reduction of its genome, the *H. elephantis* symbiont lost many metabolic capacities. However, it retained functional pathways for four B vitamins, a trait typical for symbionts in blood-feeding insects. Considering genomic, metabolic, and phylogenetic characteristics, the new symbiont closely resembles those known from several sucking lice rather than chewing lice.

**Importance:** Rhynchophthirina is a unique small group of permanent ectoparasites that is closely related to both sucking and chewing lice. These two groups of lice differ in their morphology, ecology, and feeding strategies. As a consequence of their different dietary sources, such as mammals’ blood versus vertebrate skin derivatives, they also exhibit distinct patterns of symbiosis with obligate bacterial symbionts. While Rhynchophthirina shares certain traits with sucking and chewing lice, the nature of its obligate symbiotic bacterium and its metabolic role are not known. In this study, we assemble genome of symbiotic bacterium from *Haematomyzus elphantis* (Rhynchophthirina), demonstrating its close similarity and phylogenetic proximity to several symbionts of sucking lice. The genome is highly reduced (representing the smallest genome among louse-associated symbionts) and exhibits a significant loss of metabolic pathways. However, similar to other louse symbionts, it retains essential pathways for the synthesis of several B vitamins.

## Introduction

The insect infraorder Phthiraptera includes several exclusive ectoparasitic groups with different feeding strategies. The two main types are sucking lice (monophyletic parvorder Anoplura) and the polyphyletic/paraphyletic assemblage known as ‘chewing lice’ (1). The former developed a unique arrangement of mouthparts by creating stylets that are used exclusively to feed on mammalian blood. The latter retained a more primitive type of chewing mouthparts and can utilise a broader variety of food sources (vertebrate skin products, fur, feathers, etc.). Since both these groups live on nutritionally compromised diets, they harbour symbiotic bacteria that provide them with the missing compounds, particularly B vitamins (2–4). These symbionts differ in many features from free-living bacteria. As a rule, they undergo dramatic genomic changes during their transition from a free-living bacterium to obligate symbionts, particularly manifested by a reduction in genome size, loss of many metabolic pathways, and shift of the nucleotide composition toward AT (5). The degree of these changes is determined by the duration of the symbiosis and its intimacy with the host (i.e., facultative versus obligate symbionts).

The current state of knowledge suggests that within Phthiraptera, symbionts have reached the most advanced evolutionary stages in several groups of sucking lice (Table 1 shows that the most reduced genomes with lowest GC content belong to the symbionts of Anoplura). This probably reflects the differences in diet sources between the phthirapteran groups, as symbionts with strongly reduced genomes are also found in other insects that obtain their nutrients in all stages of development exclusively from vertebrate blood, such as tsetse flies (6), hippoboscids (7), and cimicids (8). In these obligate blood feeders, the presence of the symbiont is essential. They therefore develop a direct vertical transmission of the symbionts to the progeny (usually by transovarial transfer), resulting in a strict long-term host-symbiont coevolution. This long and intimate association, in turn, leads to strong changes in genome features.

**Table 1.**
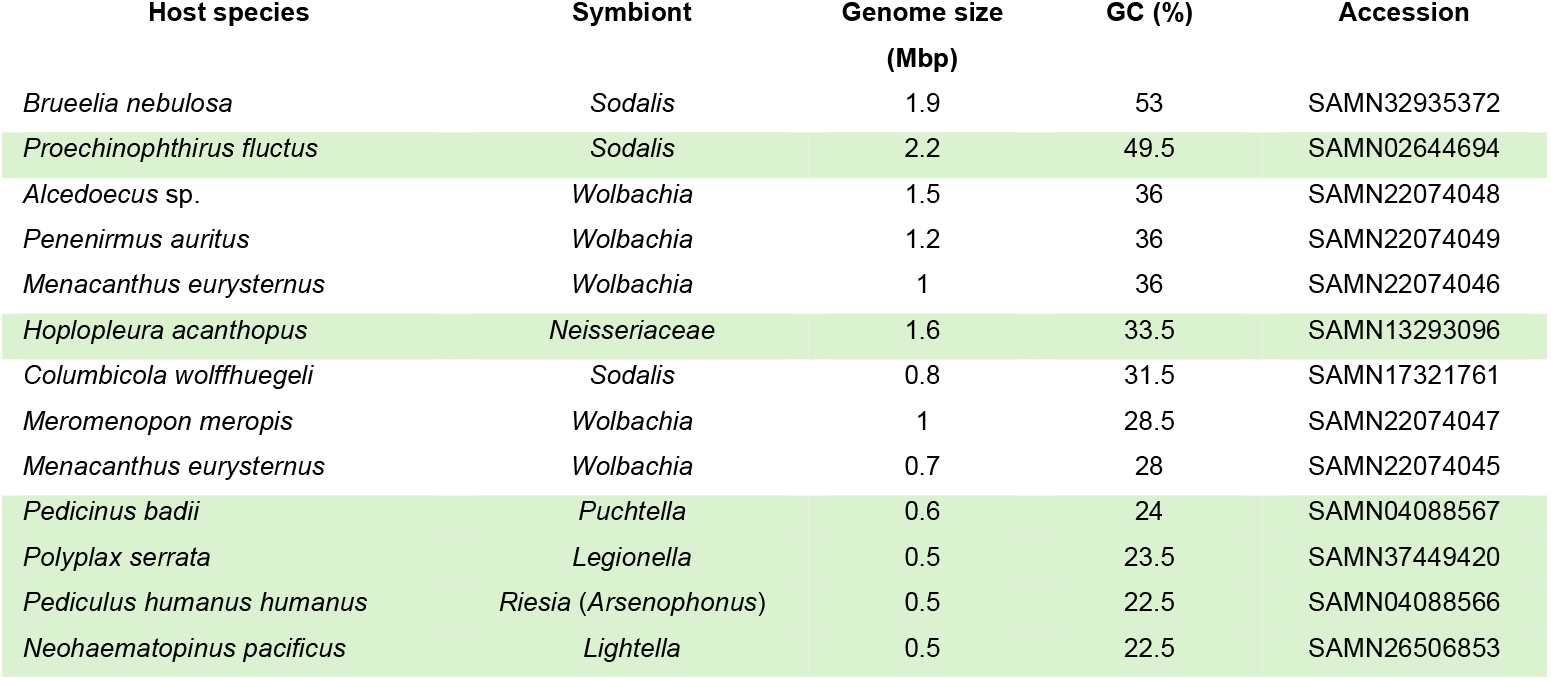
Genome sizes and GC contents of obligate symbionts described from Phthiraptera. Green highlight = symbionts of Anoplura.

Screening of several groups of Anoplura revealed a surprising diversity of symbionts, suggesting multiple acquisitions/loses/replacements of the symbionts in the evolution of course of the sucking lice (9, 10). Due to this dynamic process, sucking lice display not only a broad diversity of phylogenetic origins, but also different stages of their genomic evolution (explaining large genomes of the *Proechinophthirus* and *Hoplopleura* symbionts in Table 1). This contrasts with ‘chewing lice’, where only two bacterial genera, *Wolbachia* and *Sodalis*, were found as nutritional symbionts (3, 11, 12).

Apart from the two main types (Anoplura and ‘chewing lice’), Phthiraptera include a small group of Rhynchophthirina, with only three known species, living on elephants and African warthogs (13). Phylogenetically, they are related to Anoplura or to the ‘chewing lice’ family Trichodectidae (1). Their feeding mode and morphological adaptation differ from those of the other Phthiraptera. They possess an elongated rostrum with mandibles, which are used to feed on the host’s skin. However, it is not known whether they feed only on skin debris or also on host blood. The presence of an obligate symbiont in *Haematomyzus elephantis* has previously been indicated by a 16S rRNA gene screening (10). Comparison of this gene across free-living and symbiotic bacteria revealed that the amplified bacterial 16S rRNA gene of the *H. elephantis* sample showed an extreme decrease in GC content, even compared to the ancient obligate symbionts such as *Buchnera* or *Blochmannia*. Phylogenetic analysis based on this gene indicated that the symbiont originated within enterobacteria in a cluster of symbionts known from other insects, including sucking lice.

In this study, we address the question whether the genome of the *H. elephantis* symbiont corresponds to the strongly modified genomes of symbionts known from sucking lice and other blood feeders, as suggested by 16S rRNA gene analysis. Using metagenomic approach, we assemble a genome draft and use it to infer basic genome characteristics as signature of the evolutionary stage of the symbiont (i.e., the degree of genome reduction), an exact phylogenetic position using multi-gene analysis, and reconstruction of metabolic capacities as a base for assessing possible metabolic role of the symbiont for its host.

## Methods

### Samples preparation

The *Haematomyzus elephantis* specimen (He1 hereafter) was collected in 2021 from African elephant *Loxodonta africana* kept in ZOO Prague during the veterinary check. The sample was stored in absolute ethanol and kept at −20°C. Two steps were used to extract DNA from the sample. First, total DNA was extracted using the QIAamp DNA Micro Kit (QIAGEN) from the whole body of the specimen. Second, NEBNext Microbiome DNA Enrichment Kit (New England Biolabs) was applied to the extracted DNA to remove host eukaryotic insect DNA and increase the proportion of bacterial DNA. The quality of extracted DNA was assessed by gel electrophoresis, and the concentration measured with a Qubit High sensitivity kit.

### Metagenome sequencing and assembly

The DNA sample was sequenced on the Illumina NovaSeq6000 platform (W. M. Keck Center, University of Illinois at Urbana Champaign, Illinois, USA) using 2 × 250 paired-end reads. The quality of the raw reads was verified using FastQC (Andrews, 2010) and quality trimming was performed using the BBtools package (https://jgi.doe.gov/data-and-tools/bbtools). The resulting dataset contained 64,718,064 reads. The trimmed reads were assembled using SPAdes assembler with –meta option resulting in a metagenomic assembly of 93,937 contigs. To identify contigs of the symbiont, we employed BLASTn-based screening of the meta-assembly by genes of several bacteria. Since the study based on two genes (16S rRNA and Tu elongation factor) revealed the position of the *H. elephantis* symbiont within enterobacteria (10), we used four enterobacterial species for screening. Two of them were symbionts known from other lice, *Lightella neohaematopini* L207 (GCA_025728395.1), and *Puchtella* sp. str. PRUG (LKAS00000000); and one was a symbiont of the tsetse fly, *Wigglesworthia glossinidia* (NC_016893). Since symbiotic bacteria have as a rule reduced genomes with many missing genes, we also included a nonsymbiotic enterobacterium with a ‘normal’ genome, *Citrobacter rodentium* (NC_013716). We extracted all genes from these four genomes (6660 genes) and used them as a query in the BLASTn search (14) with the E-value set to 10.0 and the hit number to 1. This search retrieved 2,305 contigs with coverage ranging from less than 1x to 240x. The origins of these contigs were verified by BLASTn against the NCBI nt database. By this method, we finally identified four contigs of high coverage (approximately 80x – 240x; see Results and Discussion for details), which we consider a draft genome of the new obligate symbiont of *H. elephantis*.

To check for the presence of plasmids in the complete assembly, we used PlasmidFinder (Carattoli et al., 2014) with different sensitivity settings (95%, 85%, and 60%). The genome draft was annotated using PROKKA (Seemann 2014) and used in the downstream analyses (complete annotation is available in Mendeley Data under the “doi” link 10.17632/86tc28n56f.1). From the annotation, we derived the main characteristics of the new genome draft (including coding density, gene count, mobile elements, etc.) that were compared to the related genomes, (Table 2). The number of phages was analyzed with PHASTEST (15). The completeness of the genomes draft was assessed with BUSCO v4.0.6 (16).

**Table 2.**
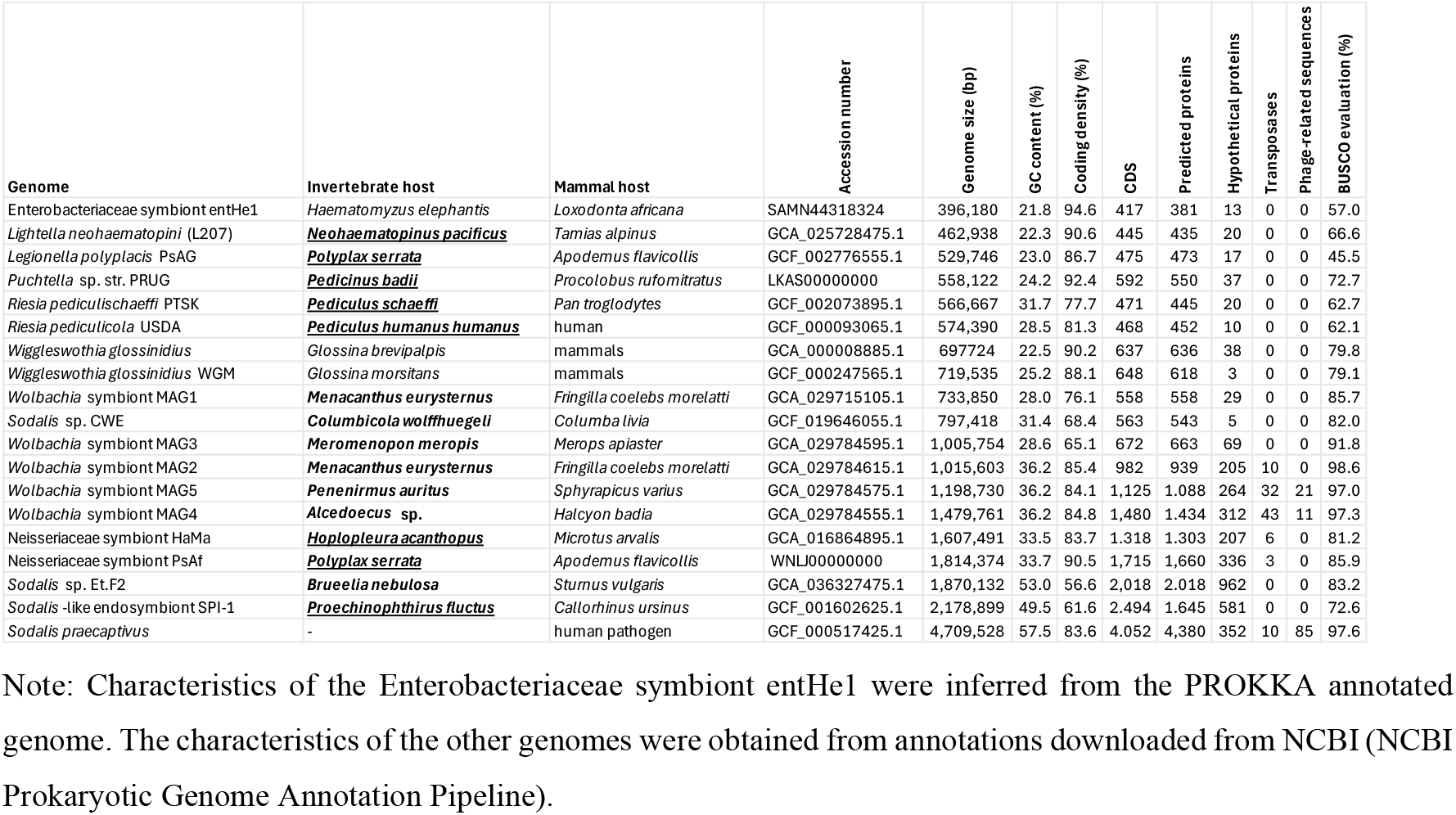
Main genome characteristics of symbionts in ectoparasitic insects. The names of sucking lice (Anoplura) are printed in bold underlined, “chewing lice” in bold.

To obtain stronger evidence that the genome belongs to an obligate symbiont, we used the published SRA data (SRR5308122) (1). We downloaded the data from ENA (https://www.ebi.ac.uk/ena/browser/home) and processed it in the same way as the reads generated from our He1 sample. The overall similarity of the two drafts was assessed by calculating average nucleotide identity in the ANI calculator (17), and by aligning the annotated contigs in Mauve (18). Taking into account the high similarity of the two genomes (ANI 99.87%; see below), only the genome draft from the He1 sample was used in the subsequent analyses.

### Phylogenetic analyses

To identify phylogenetic position of the *H. elephantis* symbiont with higher confidence than in the previous analysis based on 16S rRNA (10), we downloaded a set of 108 Enterobacteriales proteomes from the NCBI and JGI databases, including the putative closest relatives identified by BLASTp searches against the NCBI database (*Lightella, Puchtella*, and *Wigglesworthia*; accession numbers of all included proteomes are listed in sheet a of Supplementary Table S1). Several additional gammaproteobacteria representing different orders were included as outgroups: *Vibrio cholerae* O1 biovar eltor strain N16961 (Vibrionales), *Pseudomonas aeruginosa* PAO1 (Pseudomonadales), *Candidatus Evansia muelleri* CEM1.1 (Oceanospirillales), and *Xanthomonas citri* pv. vignicola strain CFBP7111 (Xanthomonadales). For this set of taxa, we extracted 14 orthologs previously identified by (19) by BLASTp search (E-value 10.0), using protein queries from *Salmonella enterica* (accession NZ_CP065718). The search retrieved sufficiently long and reliably aligned sequences from all proteomes (see sheet b of Supplementary Table S1 for the list of orthologs). The same set of orthologs was also obtained for the new Enterobacteriaceae symbiont of *H. elephantis* by performing a BLASTp search (with a default E value of 10.0) against the symbiont’s annotated genes, and the result was confirmed by visual inspection and gene annotations. The amino acid sequences of each of the 14 orthologs were aligned using MAFFT v.7.450 (20) with the E-INS-i setting in Geneious Prime v.2020.2.5 (21), visually inspected, and then concatenated. Ambiguously aligned regions were removed with Gblocks (22), applying the less stringent option. The resulting concatenated matrix (7,188 amino acids) was analysed using both maximum likelihood (ML) and Bayesian inference (BI). The ML tree was generated using the PhyML v.3.0 web server (23) under the Q.plant +G+I+F evolutionary model, identified as the best fit using the Bayesian Information Criterion (BIC) through the Smart Model Selection (SMS) algorithm (24) implemented in the PhyML web server. Considering the character of the data, which contained taxa of considerably different branch lengths and nucleotide composition, we used for the BI analysis the program PhyloBayes MPI v.1.8 (25) with CAT-GTR model to minimise this source of artifacts. The analysis was run until the maxdiff parameter dropped below 0.1 (70,000 generations). All matrices and phylogenetic trees are deposited in Mendeley Data under the “doi” link 10.17632/86tc28n56f.1.

### Reconstruction of metabolic capacity

Metabolic capacities for the symbiont of *H. elephantis* and several related bacteria (*Lightella neohaemotopini, Puchtella* sp., *Riesia pediculicola, Wigglessworthia glossinidia, Sodalis* sp., and *Sodalis praecaptivus*) as well as other obligate mutualists of lice (*Legionella polyplacis* and *Neisseriaceae* symbiont) were evaluated using the Kyoto Encyclopedia of Genes and Genomes (KEGG) server (26). Accession numbers for all compared genomes are listed in Table 2. K numbers, which link metabolic functions to the annotated genes, were identified for each genome using the BlastKOALA server (27). The K numbers for the compared genomes are deposited in Mendeley Data under the “doi” link 10.17632/86tc28n56f.1. These metabolic capacities were then mapped for each genome based on the KEGG structure of the metabolic pathways (Supplementary Table S2). From this overview and with KEGG metabolic pathways as a reference, we evaluated the completeness and potential functionality of the pathways involved in B-vitamin synthesis.

In some seemingly nonfunctional pathways (i.e. those missing single or few genes but retaining several others), we checked the possibility of compensation by the host. We used two approaches. First, we compared for the given vitamin the incomplete pathway of the symbiont with the corresponding pathway of *Pediculus humanus*, the only sucking louse with the reconstruction of the metabolic pathways available in KEGG (KEGG ID T01223). Second, we used the *P. humanus* gene as a query and screened the assembly of *H. elephantis* by BLASTn implemented in Geneious Prime with default parameters. If the search retrieved hits, we confirmed their annotation by blasting it against the nr database in NCBI. Genes and blast hits for this search are provided in Supplementary Table S3.

## Results and Discussion

### The assembly and genome characterization

The assembly of short Illumina reads produced by the SPAdes assembler contained four contigs that were assigned to a novel enterobacterial symbiont of *H. elephantis* (entHe1 hereafter). Three of these contigs had similar coverage (approximately 80x). The coverage of the fourth contig was considerably higher (approx. 240x). Since this latter contig only contained a single gene, the 23S rRNA, the coverage discrepancy is most likely due to the assembly problems when dealing with rRNA genes that occur in several copies in the genome. The presence of this single-gene contig and the terminal position of the 16S rRNA gene of one of the contigs suggests that fragmentation of the symbiont genome is caused mainly by the presence of multiple copies of the rRNA gene.

The genome of the *H. elephantis* symbiont entHe1 shows signs of a strong reduction and shift in nucleotide composition even compared to the obligate symbionts of other blood-sucking lice (Table 1). It also shares with the other strongly modified symbionts the signatures of genome economization, such as high coding density and lack of mobile elements, transposons, and phages. The completeness determined by the BUSCO analysis reached 57%. However, such low values are common for the reduced genomes of obligate symbionts (see other symbionts in Table 1; for example, *Legionella polyplacis* only reached 45.5%, although it has been reported with a close circular genome).

The assembly of the SRA data downloaded from ENA (SRR5308122) contained four contigs homological to the entHe1 genome draft. However, the overall coverage obtained with this data was significantly lower (approximately 9x – 18x, except for the 187x coverage of the 23S rRNA gene), and the draft was slightly shorter (394,551 bp vs 396,180). The size difference was due to the missing 16S rRNA gene at the terminal part of the second longest contig, obviously caused by a problem with assembling the copies of the 16S rRNA gene. Except for this difference, both drafts displayed a high similarity, with an identical number of CDSs and the average nucleotide identity 99.87%. The Mauve alignment of concatenated drafts produced a single locally collinear block with identical gene order in both genomes.

### Phylogenetic position

Phylogenetic analyses performed by maximum likelihood and Bayesian inference produced similar topologies, which recognized three clusters of long-branched obligate symbionts from various insects. According to the dominant taxa, we named the clusters i) *Buchnera* clade, ii) *Baumannia, Wigglesworthia, Blochmannia*, louse symbiont clade, and iii) *Arsenophonus*/*Riesia* clade (Figure 1). However, the two methods differed in positions of these clades. While maximum likelihood placed the *Buchnera* clade within the ‘*Baumannia, Wigglesworthia, Blochmannia*, louse symbiont clade’, the analysis based on Bayesian analysis under CAT-GTR model separated the three clades as monophyletic groups. The latter arrangement corresponds to the result obtained by the previous complex analysis of endosymbionts origins (19). It suggests that the result of the Bayesian analysis is not (at least not entirely) determined by a long-branch artifact. Within the phylogenetic tree, the obligate louse symbionts are placed into two different clusters. First, *Riesia*, the symbiont of *Pediculus* lice, branches within *Arsenophonus* clade, as demonstrated in previous studies (10, 28). Two other obligate symbionts of lice, *Puchtella* from the macaque louse *Pedicinus obtusus* (29) and *Lightella* from *Neohaematopinus pacificus* (30) are placed within a cluster rich with different symbionts, including *Wigglesworthia, Blochmannia*, and *Baumannia*. The sequence of the entHe1 symbiont was placed on an extremely long branch within this cluster, as a sister taxon to the other two louse symbionts.

**Figure 1.**
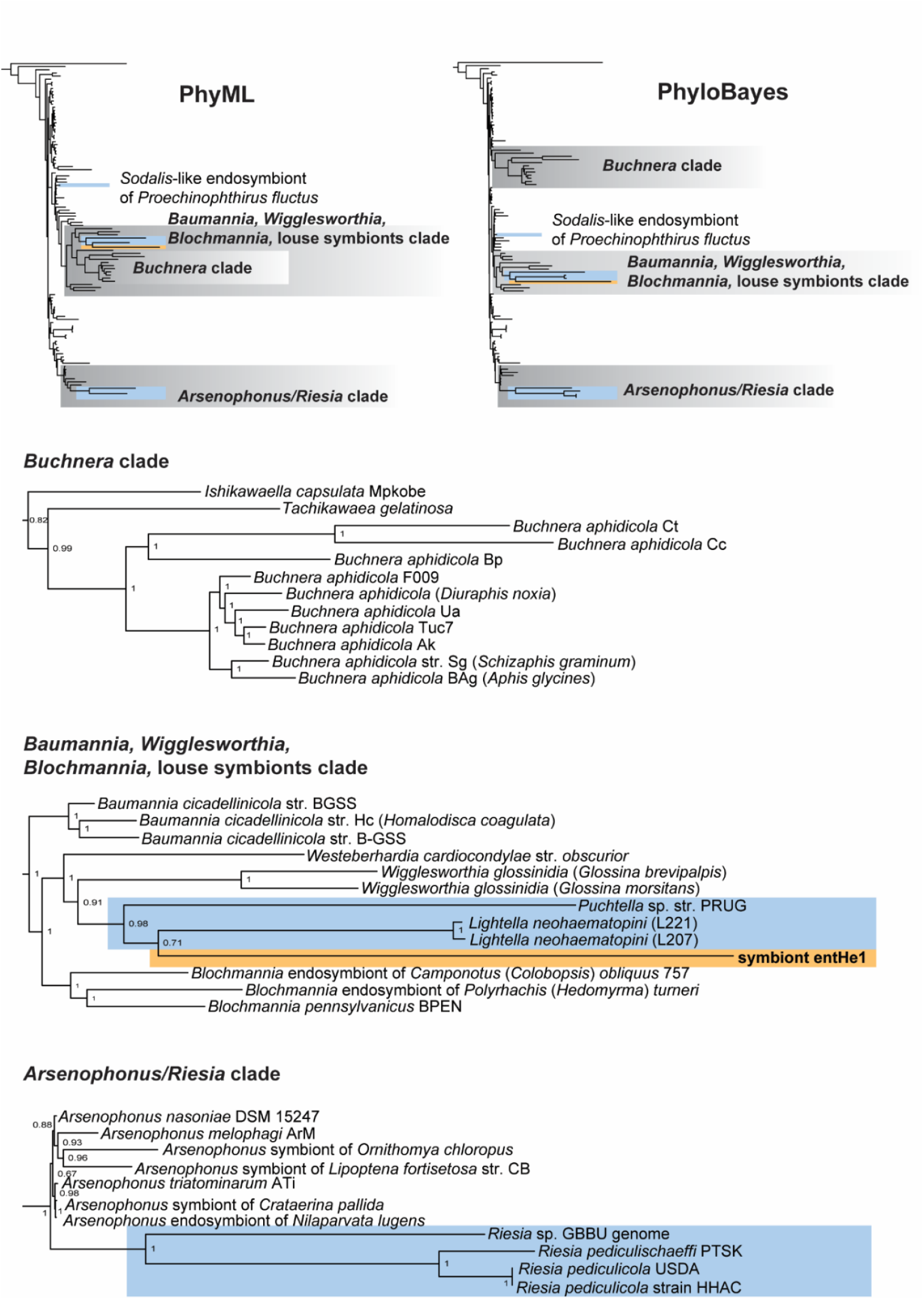
Phylogenetic trees inferred by maximum likelihood (PhyML under model Q.plant +G+I+F) and Bayesian inference (Phylobayes under model CAT GTR) from matrix of 14 genes (7,188 amino acids). The top trees show the positions of the long-branched clusters (grey background) in the analysis. The subtrees below provide details on the cluster’s arrangement in the PhyloBayes tree. Blue background = symbionts associated with sucking lice (Anoplura); orange background = new entHe1 symbiont of *Haematomyzus elephantis*.

However, it would be difficult to infer any evolutionary scenario from this clustering. The three louse genera (*Pedicinus, Neohaematopinus*, and *Haematomyzus*) are phylogenetically very distant, and their association with related symbiotic bacteria could hardly reflect any coevolutionary history. This view is further supported by the fact that symbionts of yet other louse genera belong to different phylogenetic clusters (e.g. *Riesia* in the tree in Figure 2), or even outside Enterobacteriaceae (4, 31). It should also be noted that although the Phylobayes CAT-GTR method is designed to address the problems associated with the long branches of aberrant sequences, an effect of long-branch-caused artifact cannot be ruled out entirely. Considering the extreme genomic traits of the entHe1 symbiont (long branch, strong shift of nucleotide composition, low number of genes available for the phylogeny), it is possible that the reliable traces of its evolutionary history have been erased from its genome.

**Figure 2.**
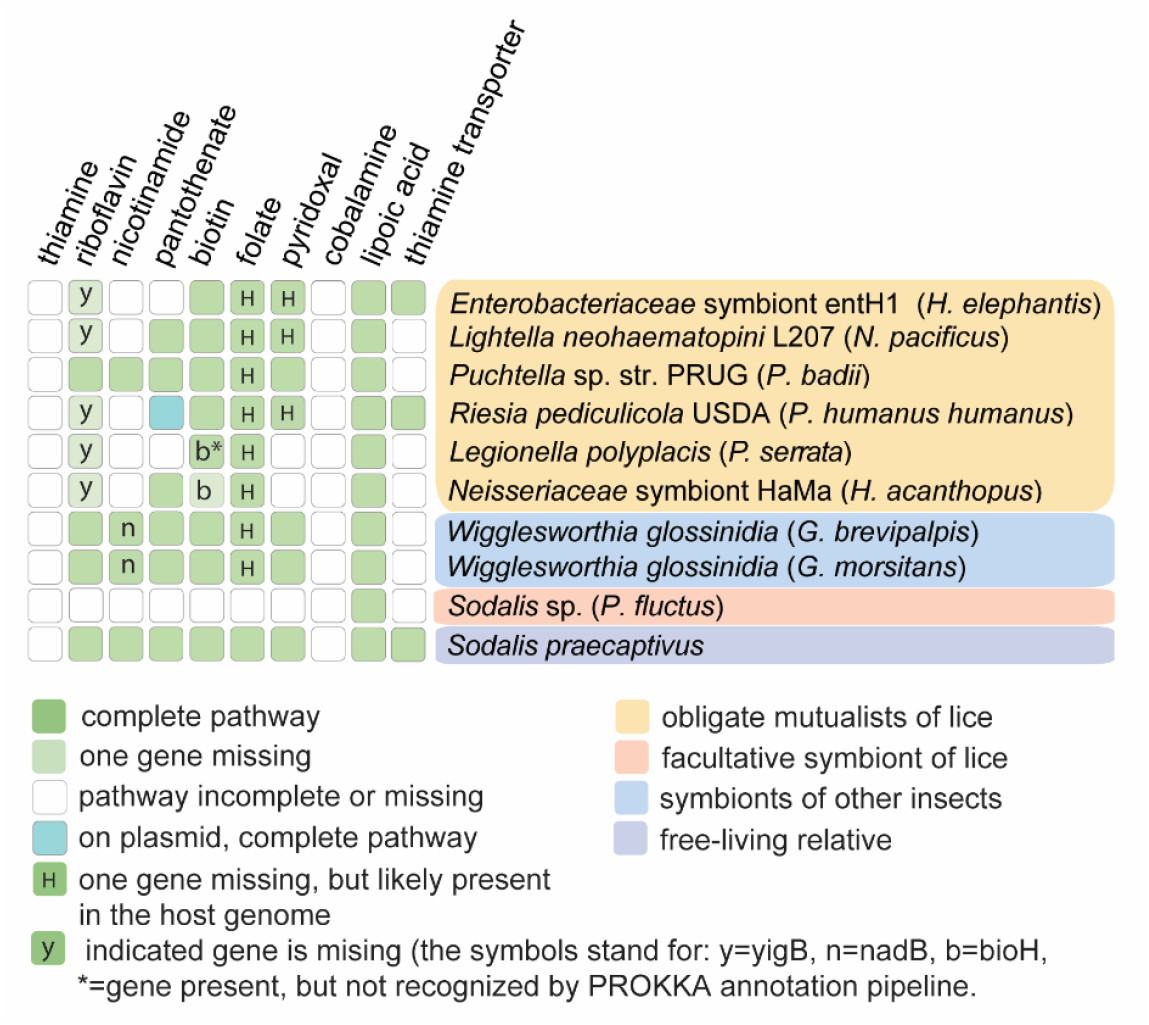
Reconstruction of metabolic capacities for vitamins B in selected bacterial genomes.

### Metabolic capacity

Reflecting its dramatic genome reduction, the symbiont entHe1 shows a significant loss/degradation of metabolic capacities, even compared to the other obligate symbionts of blood-feeding insects (Supplementary table S2). However, despite this degree of degradation, it seems to retain functional biosynthetic pathways for at least four B vitamins. This corresponds to the generally accepted view that in insects living exclusively on vertebrates blood, provision of B vitamins is the main role of their obligate bacterial symbionts.

The four vitamins are riboflavin, biotin, folate, and pyridoxal. All these pathways, except for pyridoxal, are also functional in the five remaining louse symbionts included in the analysis. Three of these pathways show identical patterns with the same missing genes in all genomes, specifically riboflavin (all louse symbiont except *Puchtella* missing *yigB*), folate (all louse symbionts missing *folE* and *phoA*) and pyridoxal (all louse symbionts missing *pdxH*). This pattern suggests that the pathways are functional despite their incompleteness, since the presence of all other genes in these pathways indicates that there is a selective pressure on the functionality of the pathway. A similar pattern has been previously reported in a comparative study of *Arsenophonus* symbionts in hippoboscids (32). In addition, closer inspection reveals that there are credible explanations for these ‘universally missing’ genes. In the riboflavin pathway, the missing yigB is responsible for a step (dephosphorylation of amino ribityl phosphate pyrimidine) which can be fulfilled by several different enzymes (33). For the folate and pyridoxal pathways, the missing genes *folE, phoA*, and *pdxH* are present in the host genome. This was confirmed by both methodological approaches (see Methods): The genes are present in the KEGG metabolic reconstruction for the louse *P. humanus*, and we found their homologue in our metagenomic assembly of *H. elephantis* He1. Such a principle, where two or even more organisms contribute to synthesis of a single compound, is well known from symbiotic systems (34).

Interestingly, the entHe1 symbiont of *H. elephantis* seems to lack the capacity for pantothenate synthesis. This contrasts with the comparative study of *Arsenophonus* obligate symbionts in hippoboscids, where pantothenate was identified as the only B vitamin whose provision by the symbionts was essential for the host (32). On the other hand, in our analysis presented here, the pantothenate pathway was non-functional not only in the entHe1 symbiont, but also in other two louse symbionts, *Lightella neohaematopini* and *Legionella polyplacis* (in *Riesia* the situation is more complex and method-dependent; see for the discussion below). This difference between *Arsenophonus* symbionts in hippoboscids (32) and our results here raise a question on the role of the symbionts in provisioning pantothenate to their blood-feeding hosts. The two studies differ in two main variables, the taxonomy (lice versus hippoboscids) and the taxonomy (Enterobacteriaceae versus *Arsenophonus*). It is therefore difficult to hypothesize if there is a connection between the taxonomy-related factors and the differences in metabolic capacities.

Moreover, reconstruction of metabolic capacities in symbionts with aberrant sequences (shift of nucleotide composition, gene truncation) may be affected by errors during gene annotation using automatic pipelines. An example shown in the analysis of (32) and relevant to this study, is the functionality of the pantothenate pathway in *Riesia*. This obligate symbiont of hominid lice *Pediculus* and *Phthirus* is considered capable of pantothenate synthesis due to the presence of core genes, *panB, panC*, and *panE* on plasmid (35). However, while the homologous sequence to the *panE* gene can be traced in all analysed *Riesia*, in some strains it is highly modified and not recognised by some of the annotation pipelines as a functional gene (32). In such cases, it is impossible to decide based on the DNA sequence alone if the gene/pathway is still functional or too degraded.

The new lineage of obligate symbiotic bacterium, reported in this study, complements an insight into the biology of the symbionts associated with different groups of Phthiraptera. The genome characteristics of this bacterium correspond to the general pattern of the obligate insect symbionts by significantly reduced genome but preserved pathways for B vitamins presumably essential for the host. The comparison shows that it closely resembles other louse associated symbionts, and even clusters together with the symbiotic bacteria known from several species of Anoplura. Although the genomic/metabolic similarity of the *H. elephantis* symbiont entHe1 with the other lice-associated bacteria is obviously due to the same evolutionary processes (relaxed selection, nutritional role), the reason for its phylogenetic proximity to several louse symbionts is less clear. As stated above, this clustering is certainly not due to a common history, i.e., the lice-bacteria coevolution. It is well known that some bacterial groups are particularly prone to establish symbiotic relationships (e.g., *Arsenophonus, Sodalis, Wolbachia*). It is interesting to consider the possibility that some of the bacteria prone to symbiosis might have affinity for a specific group of insects, such as several related louse-associated branches retrieved in our study. It will, however, require more data and perhaps further advances of phylogenetic methods to confirm if this phylogenetic arrangement is an artefact (e.g. due to an inadequate sampling or long branch attraction) or if it reflects a real evolutionary history.

## Data availability

Illumina reads were deposited in the NCBI Sequence Read Archive (SRA) repository under BioProject accession number PRJNA1173735. The genome draft of the new *Haematomyzus elephantis* symbiont entH1 is deposited in the GenBank under Accession SAMN44318324.

## Supplements

**Supplementary table S1:** List of genomes and single-copy ortholog genes used in the “14-gene matrix”.

**Supplementary table S2:** Comparison of metabolic capacities of the new Enterobacteriaceae symbiont and selected bacterial symbionts.

**Supplementary table S3** : Identification of “missing genes” in *H. elephantis* He1 genome.

